# Cycle-frequency content EEG analysis improves the assessment of respiratory-related cortical activity

**DOI:** 10.1101/2024.03.15.585182

**Authors:** Xavier Navarro-Sune, Mathieu Raux, Anna L. Hudson, Thomas Similowski, Mario Chavez

## Abstract

Time-Frequency (T-F) analysis of EEG is a common technique to characterise spectral changes in neural activity. This study explores the limitations of utilizing conventional spectral techniques in examining cyclic event-related cortical activities due to challenges, including high inter-trial variability. Introducing the Cycle-Frequency (C-F) analysis, we aim to enhance the evaluation of cycle-locked respiratory events. For synthetic EEG that mimicked cycle-locked pre-motor activity, C-F had more accurate frequency and time localization compared to conventional T-F analysis, even for a significantly reduced number of trials and a variability of breathing rhythm. Preliminary validations using real EEG data during both unloaded breathing and loaded breathing (that evokes pre-motor activity) suggest potential benefits of using the C-F method, particularly in normalizing time units to cyclic activity phases and refining baseline placement and duration. The proposed approach could provide new insights for the study of rhythmic neural activities, complementing T-F analysis.

## INTRODUCTION

Breathing is the only vital function controlled both automatically and voluntarily (Horn & Waldrop, 1998; Mohammadshirazi et al., 2023). The involvement of the brain in breathing control ranges from increased attentional resources (Moore et al., 2012), to voluntary modification of the breathing pattern (Pouget et al., 2018), as well as compensatory mechanisms in the presence of ventilatory constraints that hinder adequate airflow (Raux et al., 2007). A respiratory-related cortical activation has been described in various respiratory disorders, including chronic obstructive pulmonary disease (COPD) (Nguyen et al., 2018), obstructive sleep apnea (Launois et al., 2015), congenital central alveolar hypoventilation (Tremoureux et al., 2014) and respiratory muscle weakness (Georges et al., 2016). In this context, cortical activation is interpreted as a compensatory mechanism, and is often associated with dyspnea (defined as an upsetting or distressing awareness of breathing activity) (Manning & Mahler, 2001).

The preparation and execution of breathing movements involves specific brain areas, and both the frequency content and latencies of respiratory-related premotor and motor potentials depend on various scenarios. The generation of slow cortical (readiness) potentials in the primary motor cortex and supplementary motor area is typically observed during the preparation of self-paced breaths (Hudson et al., 2018; Macefield & Gandevia, 1991). Slow potentials occur 1.5 to 1 s before the onset of inspiration and are currently studied using averaging techniques on event-related electroencephalographic (EEG) segments (Raux et al., 2007). Pre- and post-stimuli changes in EEG activity of faster rhythms may also be present in the sensorimotor cortex. The most studied frequencies in this case are the *alpha* (8-12Hz) and *beta* (13-30Hz) bands (Pfurtscheller & Lopes da Silva, 1999), with a specific interval between 9 and 13 Hz, referred to as the *mu* rhythm, which is considered particularly insightful as it provides understanding into the neural mechanisms underlying the interaction between action and perception (Hari, 2006). Our previous investigations of respiratory-related changes in these faster rhythms had varied results (Dubois et al 2016 and Hudson et al 2016).

Changes in EEG frequencies during sensory stimulation were first quantified using the method of event-related spectral desynchronization (ERD) (Pfurtscheller, 1977). The term desynchronization, which describes an event-related decrease in activity, was later complemented with the opposite effect, that is, an event-related power increase, to describe EEG changes under the term event-related desynchronization/synchronization (ERDS) (Pfurtscheller, 1992). This technique relies on averaging a number of EEG segments, time-locked to an event (trials), and measuring amplitude changes in both the temporal and spectral domains, with respect to a reference period of a few seconds before the event occurs (baseline) (Pfurtscheller & Lopes da Silva, 1999). With time-frequency (T-F) transformations of the EEG signals, event-related changes can be represented in a spectrogram, a 2D representation that allows localization of the time-frequency components. This technique is commonly referred to as event-related spectral perturbation (ERSP) (Makeig et al., 2004).

At present, ERSP is one the most popular representations in computational neurosciences, as it allows for easy visualization of power spectral changes induced or evoked by events related to both external (visual, auditory, tactile) and internal (self-paced movement) stimuli. Current protocols typically define a fixed value of latencies and duration for each stimulus to facilitate data segmentation into trials and analyses. However, in case of self-initiated or self-paced stimuli, the interpretation of ERSP can be biased by inter-trial variability, which refers to different trial segments’ duration mainly due to fatigue or uncontrolled cognitive processes of the subject during the experiment (Fox et al., 2007; Provost et al., 2018). Although some strategies can minimize this variability (e.g. discarding short trials), the conventional procedure for computing ERSP is rarely questioned.

Several alternatives to the standard procedure have been proposed to improve the ERSP estimation. Lemm and colleagues (Lemm et al., 2009) introduced a new computational approach that measures ERSP using a dynamic reference, which considers the dynamics of unperturbed EEG rhythms instead of a constant baseline level, resulting in more accurate estimations of the evoked somatosensory activity. Another study reported that optimal ERSP detection depends on the identification of participant-specific reactive bands, and used a band-limited multiple Fourier linear combiner to demonstrate that the classification performance of cue-based motor imagery tasks was significantly increased than that of conventional ERSP estimation (Wang et al., 2012). While both methods can be useful in reducing bias in ERSP estimation under external stimuli, addressing the variability of trials related to endogenous (self-initiated), unbalanced stimuli remains a challenge for motor imagery tasks (Daeglau et al., 2020) and gait studies (Delval et al., 2020). To the best of our knowledge, the relevance of conventional ERSP for the particular case of respiratory-related cortical activity has not yet been addressed.

Breathing is controlled by automatic central pattern generators located in the brainstem and by and voluntary control pathways, both of which are in constant interaction during wakefulness (Laviolette et al., 2013; Straus et al., 2004). This can lead to high inter-trial variability in terms of segment duration and neural dynamics under certain conditions (Datta et al., 1991; Homma & Masaoka, 2008). Therefore, conventional event-related spectral analysis may not be accurate in identifying respiratory-related evoked potentials. Due to the cyclic and pseudo-periodic nature of breathing, previous studies have explored respiratory-phase domain analysis techniques. In heart rate variability (HRV) analysis, this approach provides a cycle-by-cycle view of HRV, offering advantages over time-domain techniques in quantifying respiratory sinus arrhythmia (Kotani et al., 2007; Numata et al., 2013, 2021).

In this paper, we introduce the concept of the respiratory-phase domain for analyzing respiratory-related cortical activity. First, we discuss the standard methodology used to compute ERSPs and its limitations when it comes to the specificity of experiments assessing respiratory-related cortical activity. Next, we present the cycle-frequency (C-F) representation, an approach inspired by phase domain analysis that considers the cyclic dynamics of breathing to improve the estimation of ERSP. This method is validated using a model that generates respiratory-related potentials in an EEG-like signal. Finally, we apply the cycle-frequency transformation to real EEG data, comparing and discussing the benefits and limitations of this new representation.

## METHODS

### Conventional procedure to obtain ERSPs

As for most experimental paradigms in cognitive neuroscience, the study of spectral disturbances related to respiratory events can be studied with conventional time-frequency representations. The usual procedure for obtaining these time-frequency maps (Bullock, 1992) can be summarized as follows: (i) the EEG signal is divided into *n* excerpts ***x*** = {*x*_1_, *x*_2_,…, *x*_*n*_}, of *l* seconds (*m* samples) around the events of interest; (ii) each EEG segment is transformed into the time-frequency domain, producing a complex variable ***X*** = {*X*_1_, *X*_2_,…, *X*_*n*_} with *X*, ∈ ℂ*^m^*^×*q*^, where *q* denotes the number of frequencies; (iii) the absolute values of *X_i_* are averaged to obtain a single image representing the power of induced activity, 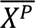, whereas the complex values of *X_i_* are used to compute the phase alignment between trials, often referred as intertrial coherence, 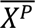; (iv) finally, average maps are normalized with respect to a baseline period. Evoked activities can be similarly obtained by averaging time-frequency transformation *X_i_* of all epochs, and by taking the absolute value of the final map.

To identify the regions with relevant power changes, statistical significance can be assessed by cluster-based permutation tests (Bullock, 1992) or bootstrapping (Gross et al., 2013). However, statistical tests are performed on grand-average maps, which are obtained by averaging an ensemble of maps from individual participants. The results of standard ERSP analysis strongly depend on the choice of:

1. The length of the segments (*l*), which is fixed according to the experimental conditions. The segments are centered around the event of interest (*t_0_* = 0 s) and divided into a pre-event period (*t_pre_*) and a post-event period (*t_post_*). The length of each period should be chosen to capture the potentials of interest while avoiding overlap with neighboring trials.
2. The baseline period, which should be distant from the onset of the potentials of interest to ensure a good contrast. Typically, it is a time interval extending up to 20-25% of *t_pre_* or *t_post_*, depending on the study (Picton et al., 2000). Various techniques can be used to correct by the baseline, but in general, the activity of the normalized maps is reported as the average power per frequency of this period. These changes in T-F are represented as percentage, standard deviation, or decibels.

Although the conventional procedure to obtain ERSPs can be useful for the estimation of respiratory-related potentials under ventilatory constraints (Dubois et al., 2016; Hudson et al., 2016) it has significant limitations due to the inter-trial variability caused by different duration length of respiratory cycles (see Figure 1). Indeed, the resting respiratory activity in humans is characterized by its breath-by-breath variability (van den Bosch et al., 2021), which stems from the intrinsic complexity of brainstem automatic neural command (Mangin et al., 2008) and from suprapontine perturbations. This variability pertains to tidal volume (the volume inhaled during a single breath), the respiratory cycle period, the duration of inspiration, and the duration of expiration (Fiamma et al., 2007). It can result in coefficients of variation between 15 and 40% for these variables (Fiamma et al., 2007), which limits the use of time-frequency maps.

**Figure 1:**
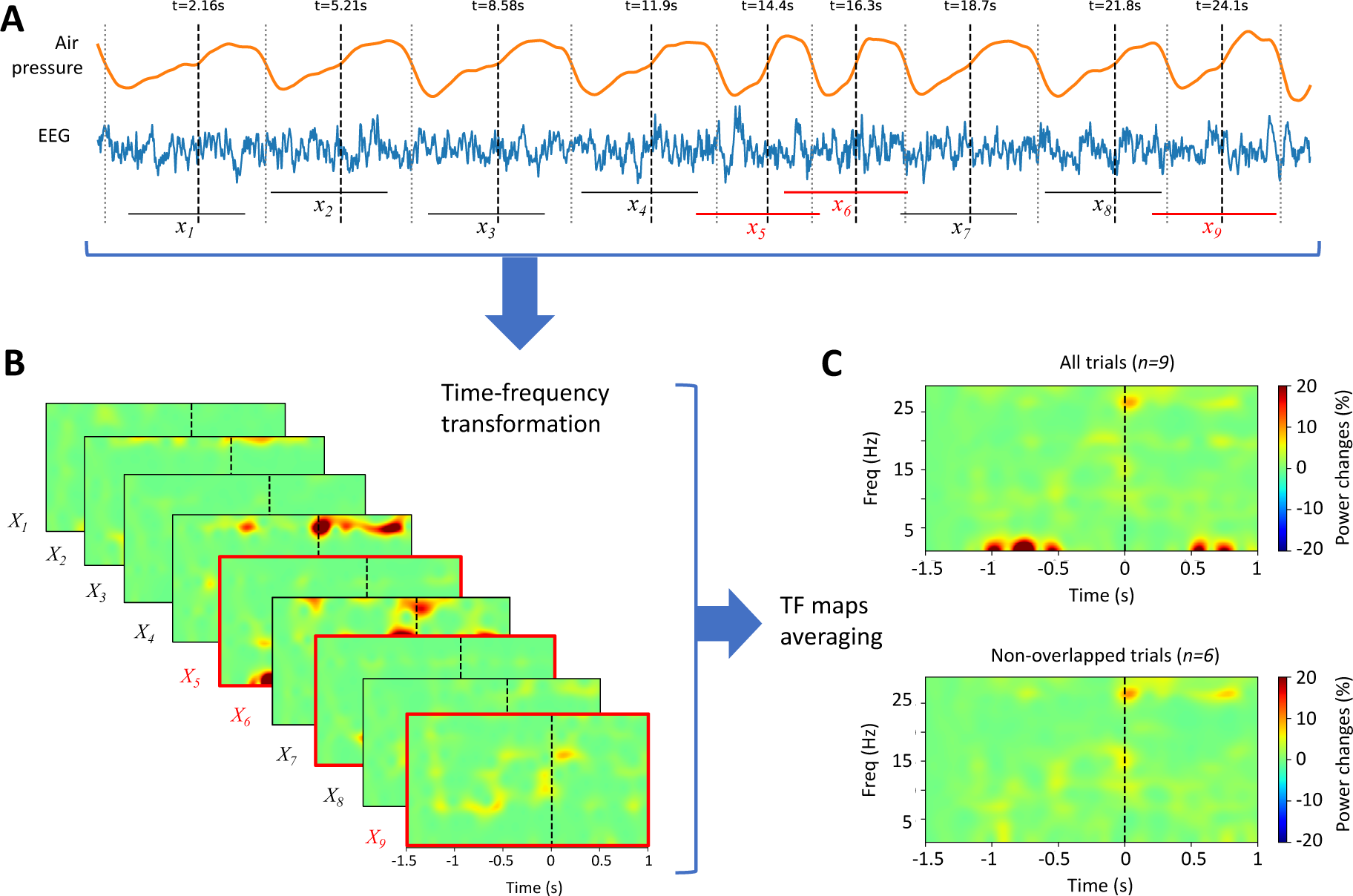
Illustration of the conventional procedure for computing ERSPs to study pre-inspiratory potentials using EEG recorded during spontaneous breathing. A: The EEG signal is divided into nine trials (x_1_ to x_9_) according to the onset of inspiration (black vertical dashed lines), and pre-inspiratory and post-inspiratory times (in this example t_pre_=-1.5s and t_post_=1s). The onset of expiration is marked with grey vertical dotted lines and trials with significant overlap (i.e. when the next trial commences prior to 90% completion of the current trial) are highlighted in red. B: For each trial, time-frequency transformation (using complex Morlet wavelet) and baseline correction (between −1.5 and −1.25 s) are performed, resulting in n=9 time-frequency maps (X_1_ to X_9_). C: The average power maps are obtained by averaging all trials (upper map) or using only non-overlapping trials (lower map).

### Alternative procedure to obtain ERSPs: the hypothesis of cycle-dependent potentials

Several studies suggest that respiratory-related potentials may be dependent on the phase of the breathing cycle. For instance, Davenport et al (1986) found that the latencies of respiratory-related somatosensory evoked potentials depend on the intensity of the underlying neural drive to breathe, and hypothesized that somatosensory potentials induced by loaded breathing are delayed in comparison to analogous evoked potentials for electrically evoked somatosensory potentials, due to the time required for the stimulus to activate sufficient inspiratory afferents to evoke a response (Davenport et al., 1986). A recent study demonstrated that intracortical EEG and the breathing signal are significantly synchronized during volitional breathing by using spectral coherence techniques (Herrero et al., 2018; Watanabe et al., 2023).

Based on these studies, we hypothesize that breathing potentials, which typically exhibit a non-regular cyclic pattern, should be considered “cycle-locked” rather than time-locked potentials. Regarding the reference event (inspiration), pre- and post-inspiratory potentials can be distinguished. Although they may have different implications for the processing of somatosensory or (pre-)motor information, we will refer to them as inspiratory potentials for simplicity. This assumption has two practical implications for our analysis: i) inspiratory potentials start at a specific phase of the breathing cycle; hence their latencies are jittered in time according to the cycle duration and; ii) inspiratory potentials are active during a specific phase interval, and their duration is proportional to the cycle length.

To represent ERSPs using the phase of the respiratory cycle rather than time, we propose to construct cycle-frequency maps. While approximating each breath as a whole cycle (2*π* radians) between marks may not perfectly reflect the different durations of inspiration and expiration, this choice offers advantages for our study:

- Practicality: A symmetrical cycle simplifies the analysis and reduces computational complexity compared to modeling variable inspiration/expiration times.
- Focus on Inspiration: Our primary interest lies in pre-inspiratory potentials, which are likely limited to a specific window within the breathing cycle, as supported by the studies mentioned in the introduction. These studies suggest that pre-motor activity synchronizes with the inspiratory trigger, implying the oscillations of interest primarily occur during the expiration time of the preceding period.

Therefore, we define each breath as a complete cycle, with the phase of the cycle (ϕ) set to π radians at the expiration onset (similar to using inspiratory marks).

To segment the EEG, we define the analysis phase interval [ϕ_5_, ϕ_6_]. For pre-inspiratory potentials, the interval could include half of the previous breath (i.e., from the expiration onset at ϕ_5_ = −π rad) and a portion of the inspiration (e.g. ϕ_6_ = π/2 rad). Similar to T-F analysis, the events of interest (inspiration marks) are placed at ϕ = 0. Importantly, this cycle-based approach systematically avoids the inclusion of the entire previous inspiration period (ϕ < −π rad), which might also contain pre-inspiratory-related cortical activities. This ensures accurate baseline placement by selecting the 15-20% leftmost portion of the pre-inspiratory time within the current cycle (see Figure 2). If the analysis concerns post-inspiratory potentials, the phase interval can be modified to center the window to the activity of interest.

**Figure 2:**
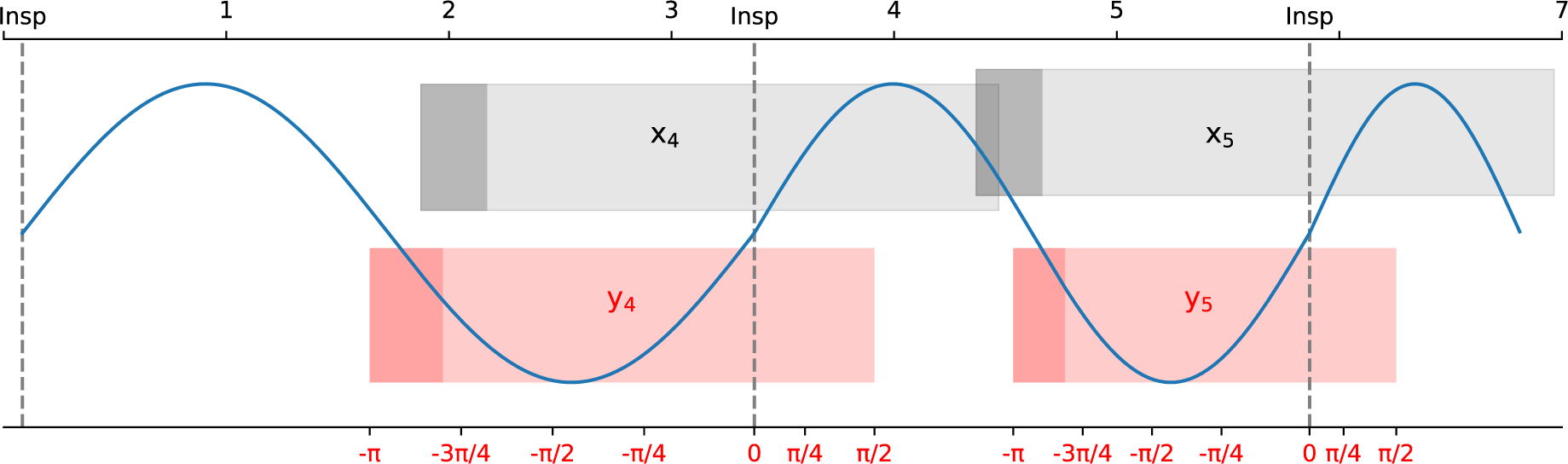
Illustration of two segmentations adapted to cycles of different duration. The breathing trace is approximated by a quasi-sinusoidal period between consecutive inspiratory marks. Grey upper rectangles correspond to conventional trials (x_4_ and x_5_) of equal length (l_4_=l_5_=2.5s), with fixed baselines (dark grey areas). Red rectangles correspond to cycle-segmented trials (y_4_ and y_5_) of variable length with a phase interval [-π, π/2] rad. Dark red areas correspond to the baseline periods adapted to each cycle by taking the 20% of the pre-inspiratory time, equivalent to the phase interval 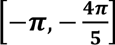

#### Construction of cycle-frequency maps

To construct cycle-frequency maps, we start by adaptively segmenting EEG as described above. Cycle-based segmentation yields *n* segments, denoted as ***y*** = {*y*, *y*_2_,…, *y_n_*}, with *y_i_* of variable length *l_i_* in seconds, (*i* = 1,2,…*n*) containing a number of samples *m_i_* = *l_i_f_s_*, with *f_s_* denoting the sampling frequency. We denote the shortest and longest breaths by *l_min_* and *l_max_*, respectively.

The next step is to compute the T-F representation on ***y***, choosing the optimal transformation parameters. As mentioned earlier, there are multiple decomposition approaches, but the most common is based on the continuous wavelet transform (Cohen, 2014). This T-F representation convolves the EEG signal with a set of complex Morlet wavelets, defined as complex sine waves tapered by a Gaussian (Tallon-Baudry & Bertrand, 1999) as follows:

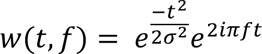

where *i* is the imaginary unit given by 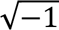 and σ = η/2πf is the width of the Gaussian. The parameter η, also referred as to the number of cycles, determines the time-frequency precision trade-off. A smaller η results in higher temporal resolution but lower frequency resolution, whereas a larger η results in higher frequency resolution but lower temporal resolution. Although η can be constant, in neural time-series it is recommended to use a variable number of cycles as a function of frequency, starting from a few cycles for the lowest frequency and increasing up to several cycles for the highest frequency of interest (Cohen, 2014).

The choice of parameters for computing the T-F representation includes the boundary frequencies represented in the spectrograms (*f_min_*and *f_max_*), and the frequency bin (*f_b_*), which define the number of frequencies (*q*). The choice of *f_min_* is generally determined by the shortest trial. Following the recommendations (Cohen, 2014), at least three complete oscillations of the lowest frequency to be analyzed should be contained in the signal. Therefore, we set *f_min_* ≥ 3/*l_min_*. On the other hand, the highest frequency is limited only by the Nyquist frequency, hence *f_max_* ≤ *f_s_*./2 even if, in practice, *f_max_*can be set to a much lower value, where most relevant information in EEG is contained.

The time-frequency transformation of *y* will produce the corresponding spectrograms, ***Y***′ = {*Y*′_1_, *Y*′_2_,…, *Y*′*_n_*}, with *Y*′*_i_* ∈ ℂ^m^_i_^×q^, where *m_i_* denotes the number of samples of trial *i*. Next, ***Y’*** is converted from time to cycle domain to obtain ***Y*** = {*Y*_1_, *Y*_2_,…, *Y_n_*}, with *Y*, ∈ ℂ*^p^*^×*q*^. The number of phase values (bins), *p,* must satisfy the condition 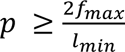 to ensure the correct representation of the highest frequencies. Likewise, the phase interval of interest Δϕ = ϕ*_e_* – ϕ*_s_* have to be taken into consideration. For instance, given a set of T-F maps with *l_min_* = 1.5 s, *f_max_*=30 Hz and Δϕ = 3π/2 rads (or 270°), choosing a phase bin *r_ϕ_* = π/90 rads (or 2°) results in *p* = Δϕ/*r*_ϕ_ = 135, which satisfies the condition 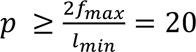.

Once the value of *p* is properly selected, the conversion from time to cycle domain can be achieved by downsampling by a trial-dependent factor, 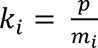. To minimise the aliasing effect of downsampling, we employed here a poliphase filter (Hsiao, 1987). Finally, a baseline correction is applied by averaging *Y*, to obtain the power, 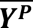, and to compute inter-trial phase coherence from the complex values *Y_i_*

### Validation strategy

To validate the proposed method, we first used artificial EEG-like signals containing cycle-locked activity of known frequency and amplitude, specifically designed to study pre-inspiratory potentials. Then, we applied the conventional and the cycle-frequency procedure to obtain ERSP and compared them. Subsequently, we tested both methods on real EEG data obtained from healthy participants breathing under different respiratory conditions.

#### Synthetic data generation

We generated artificial EEG signals using two distinct generative models: the first model incorporates pre-inspiratory activity in accordance with the hypothesis of cycle-locked potentials while the second model generates time-locked potentials to inspiratory marks. In both models, artificial EEG was generated by combining background activity based on Gaussian noise with superimposed sinusoidal bursts. These bursts mimic synchronization of EEG activity seen with movement.

For the cycle-locked model, the onset of each burst was determined by the phase of the cycle (set to 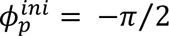), and its duration was proportional to the cycle length (burst phase interval, Δϕ*_p_* = π/4). For the time-locked model, the onset latency of oscillations was within the typical range of 300 to 700 milliseconds prior to trigger marks (Kilavik et al., 2013) and their duration was arbitrarily set to half the value of the latency.

The frequency of the sinusoidal bursts (*f_p_*) was set to 20 Hz for both models, a value typically found in pre-motor activity (Kilavik et al., 2013). Bursts were tapered by a Hanning window of the same duration to smooth the edges. The background noise follows a Gaussian distribution rescaled by a factor of 10^-5^, and the sinusoidal bursts peak was set to 5 μV. Given the targeted activity, the frequencies of interest for time-frequency transformations were *f_min_*=5 and *f_max_*=30 Hz with steps of 0.5 Hz, hence *q=50*.

To create respiratory event-related EEG epochs, we generated pseudo-inspiratory markers reflecting the typical human breathing range (2-6 seconds per breath) using a log-normal distribution (Suki, 2002; P. D. Wagner et al., 1974). A sequence of N log-normally distributed random numbers of mean μ_r_ = 4 seconds and variance σ_r_ provided the periods of the simulated breathing rhythm. The log-normal distribution controlled inter-trial variability. Higher variance (σ_r_ > 1) yielded a more dispersed distribution, reflecting greater variability in breathing cycles. Conversely, a variance close to 0 produced a skewed distribution, mimicking a more constant breathing rate. Only cycles exceeding 2 seconds, corresponding to realistic breathing patterns, were retained.

Using this approach, we created synthetic EEG at *f_s_*=500 Hz simulating breathing constraints that induced pre-inspiratory potentials cycle-locked and time-locked to the inspiration marks, with a mean respiratory period, 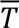, of 4 s.

To compare conventional time-frequency to cycle-frequency representations, we obtained three sets of segments for each one of the generative models:

1. ***x*** was the result of performing conventional fixed length segmentation, *l=2* s (*t_pre_*=-1.5 s; *t_post_*=0.5 s). The number of samples per trial, *m*, was 1000.
2. ***y*** was composed of mean-adapted segments, i.e. excerpts segmented according to the mean duration of breaths (4s). The phase span was 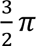 radians with ϕ*_i_* = −π and 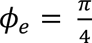 but segments had fixed length, *l=3 s* (*t_pre_*=-2 s; *t_post_*=1 s) and *m=*1500.
3. ***z*** was composed by cycle-adapted segments, with *l_i_* depending of the duration of each breath. The phase span was 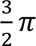 radians with ϕ*_i_* = −π and 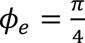.

In the following sections, each method to compute the different representations will be denoted as:

- Method A: Conventional procedure to obtain time-frequency maps from fixed lengh segments (***x***)
- Method B: Cycle-frequency transformation using mean-adapted segments (***y***)
- Method C: Cycle-frequency transformation using cycle-by-cycle adapted segments (***z***)

#### Quantification metrics for synthetic data

To assess quantitatively the efectiveness of the proposed method in handling inter-trial variability and its impact on the significance of respiratory-related potentials, we employed permutation testing (Bullock, 1992), a non-parametrical statistical approach commonly used in ERSP analysis. Briefly, the permutation testing procedure is as follows: i) Collect time-frequency maps either from two different conditions, or from a single condition to test significance with respect to a baseline period ii) Calculate the observed F-statistic, which compares the variance between the conditions to the variance within each condition, for each time-frequency point iii) Combine the data, shuffle the condition labels, and recalculate the F-statistic for many permutations to create a null distribution iv) Compare the observed F-statistic to this distribution to determine p-values and assess significance.

While the number of significant values (time-frequency points) provides an indication of the extent of significant regions, the F-statistic values offer additional insight into the strength of this significant differences. Hence, by comparing the average values of F-statistics in time- and cycle-frequency representations, we assessed:

-The number of trials: since ERSPs are obtained by averaging a number of trials, the signal-to-noise ratio generally increases with the number of repetitions. Hence we generated a variable number of trials (N = [10,…100]) and compared F-statistics with the same level of respiratory variability (σ_r_ = 1).
-The inter-trial variability robustness: by changing the degree of inter-trial variability (governed by σ_r_), we can assess the robustness of time- and cycle-frequency representations.

### Tests on real data

To illustrate the potential of our proposal, we compared qualitatively standard Time-Frequency (T-F) and Cycle-Frequency (C-F) maps using real data from healthy subjects who breathed under two distinct conditions: unloaded breathing (UB) and loaded breathing (LB) (Hudson et al., 2022). During the experiment, EEG data (32-channel system following the International 10-20 System) was recorded along breathing signals (air pressure and airflow) and accelerometry of the head. In the LB condition, subjects experienced increased effort to maintain adequate airflow due to the application of an inspiratory threshold load, whereas in UB, subjects breathed normally. We chose these data as in LB, we expect pre- and post-inspiratory potentials to arise not only due to ventilatory compensation (pre-motor and motor activity) but also due to the unpleasant sensations associated with breathing difficulty (somatosensory activity).

Time-frequency and cycle-frequency analyses were conducted using the same parameters as for synthetic signals. After obtaining different maps for each channel, we computed an average map covering the pre-motor area (average of positions F3, Fz, F4, FC1, and FC2) and an average map covering somatosensory positions (average of C3, Cz, C4, CP1, and CP2) to investigate differences between the three methods. These areas are of particular interest in the analysis of respiratory-related potentials as they reflect the preparation and execution of breaths and the sensory perception of breathing.

We also quantified the relative mu and beta power bands before the inspiratory onset. To do this, we computed the power in these bands by channel and averaged the values across all trials. The studied channels were positioned at frontal (F3, F4, F7, F8, Fz), fronto-central (FC1, FC2, FC5, FC6), central (C3, C4, Cz), centro-parietal (CP1, CP2, CP5, CP6) and parietal (P3, P4, P7, P8, Pz) cortices as they encompass movement-related activity. Values obtained in the two conditions (UB and LB) were compared using a Wilcoxon signed-rank test

#### Quantification metrics for real data

Quantifying the reliability of the different methods in real data presents several challenges. One key difficulty lies in separating the influence of confounding factors due to protocol design, noise and fluctuations within a subject. Additionally, the relative dominance of cycle-locked potentials versus time-locked potentials is unknown. This overlap makes it difficult to definitively isolate the specific contribution of each method.

Nevertheless, we propose to employ mu and beta band power as effectiveness metric (Neuper & Pfurtscheller, 2001). They are typically measured between 8-13 Hz and 14-30 Hz, respectively. Higher mu and/or beta power is generally associated with decreased cortical activation, particularly in the pre-motor and sensorimotor cortices. In particular, a decrease in mu power, or mu suppression, is often observed prior to movement initiation and during motor imagery, suggesting a role in motor preparation and planning (Hallett, 1994; Sanes & Donoghue, 2000).

The latencies of interest for both bands were set at [-pi/4, 0] radians for cycle-locked potentials and [−0.5, 0] seconds for time-locked potentials, an equivalent area in cycle- and time-frequency representations.

## RESULTS

### Tests on synthetic data

The time-frequency transform of synthetic signals was obtained by means of a Morlet wavelet which number of cycles η increases with frequency. Thus, the frequencies of the wavelets ranged from 2.5 Hz to 14.75 Hz in 50 frequency bins. This resulted in ***X***, with ***X***_***i***_ ∈ ℂ^1000×50^. For cycle-frequency maps, a phase bin **r**_ϕ_ = 1° (with a *p=*270) satisfied the restriction imposed by *l_min_*=2 s and *f_max_* = 30 Hz. Therefore, the cycle-frequency transformation of ***y*** and ***z*** resulted, respectively, in ***Y*** and ***Z***, with ***Y***_***i***_, ***Z***_***i***_ ∈ ℂ^270×50^. Table A1 (see Appendix section) summarizes all cycle-frequency parameters and the justification for the chosen values.

All power maps were baseline-corrected the leftmost 15% of samples to visualize power ratio changes. Data generated using the cycle-locked and time-locked models are denoted, respectively, with superscripts (1) and (2). Standard permutation tests were conducted to compare the significance of pre-inspiratory activity against the baseline period. Tests involved 1000 permutations and a significance level of 0.05 with threshold-free cluster enhancement (Smith & Nichols, 2009). All computations were performed using Python 3.7 language with the MNE package (Gramfort et al., 2013).

Figure 3 shows time- and cycle-frequency transformations of artifical data generated by the cycle-locked model at a mean respiratory rate μ_r_ = 4s and respiratory variability modulated by σ_r_= 1. As shown, cycle-locked potentials appeared around 20Hz, but with varying power values and time precision. Using method C (our cycle-related method, based on averaging **Z*^(1)^***), bursts (i.e. EEG synchronisation) were more noticeable and well identified in phase, whereas their temporal localization in maps computed by method A (the average of ***X^(1)^***) spaned over 0.5 seconds and the borders were not well-defined. We also observed that using method B (segmentation based in the mean duration of breaths, ***Y^(1)^***) we can identify the cycle-locked potentials, although with lower power values and precision compared to method C.

**Figure 3:**
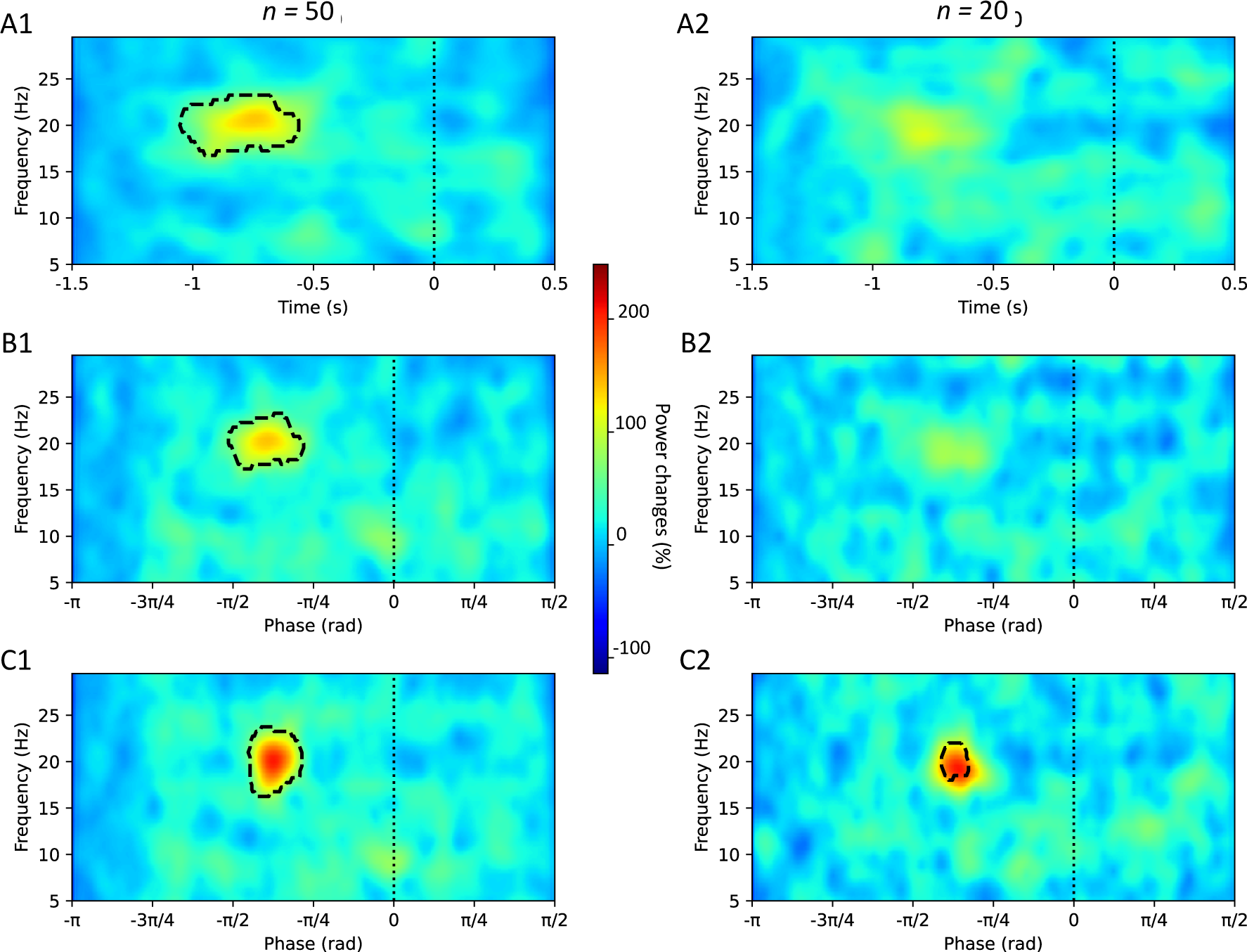
Cycle-frequency maps derived from synthetic EEG generated by the cycle-locked model. Rows A, B and C illustrate different power representations: Row A shows the map obtained by method A (conventional time-frequency transformation of fixed-length segments, ***X***^(1)^). Row B presents the map computed by method B (cycle-frequency transformation employing segments with a fixed duration based on the mean respiratory rate, ***Y***^(1)^). Row C displays the map obtained by method C (cycle-frequency transformation using cycle-adapted segmentation, ***Z***^(1)^). Left plots (A1, B1, C1) depict results from n=50 simulted trials, while right plots (A2, B2, C2) show equivalent representations from another simulation with n=20 trials. The color scale expresses the percentage of power changes with respect to the baseline. Black dotted lines enclose statistically significant areas found by the permutation test.

Permutation tests confirmed that only power maps obtained from method C retained a good precision to localize cycle-related oscillations, even when the number of averaged trials decreased from *n*=50 to *n*=20 (see Figure 3-C2). As expected, the in-phase alignment of cycle-locked activities maximized the signal-to-noise ratio.

Figure 4 illustrates time- and cycle-frequency transformations of artifical data generated by the time-locked model. Conventional time-frequency transformations effectively detected time-locked potentials, with statistically significant regions aligning with the latencies and frequency of the bursts, observerd consistently across simulations with n=50 and n=20 trials (panels A1 and A2). However, maps using method B revealed reduced significant areas corresponding to the bursts due to time-to-cycle conversion losses, as evident in panels B1 and B2.

**Figure 4:**
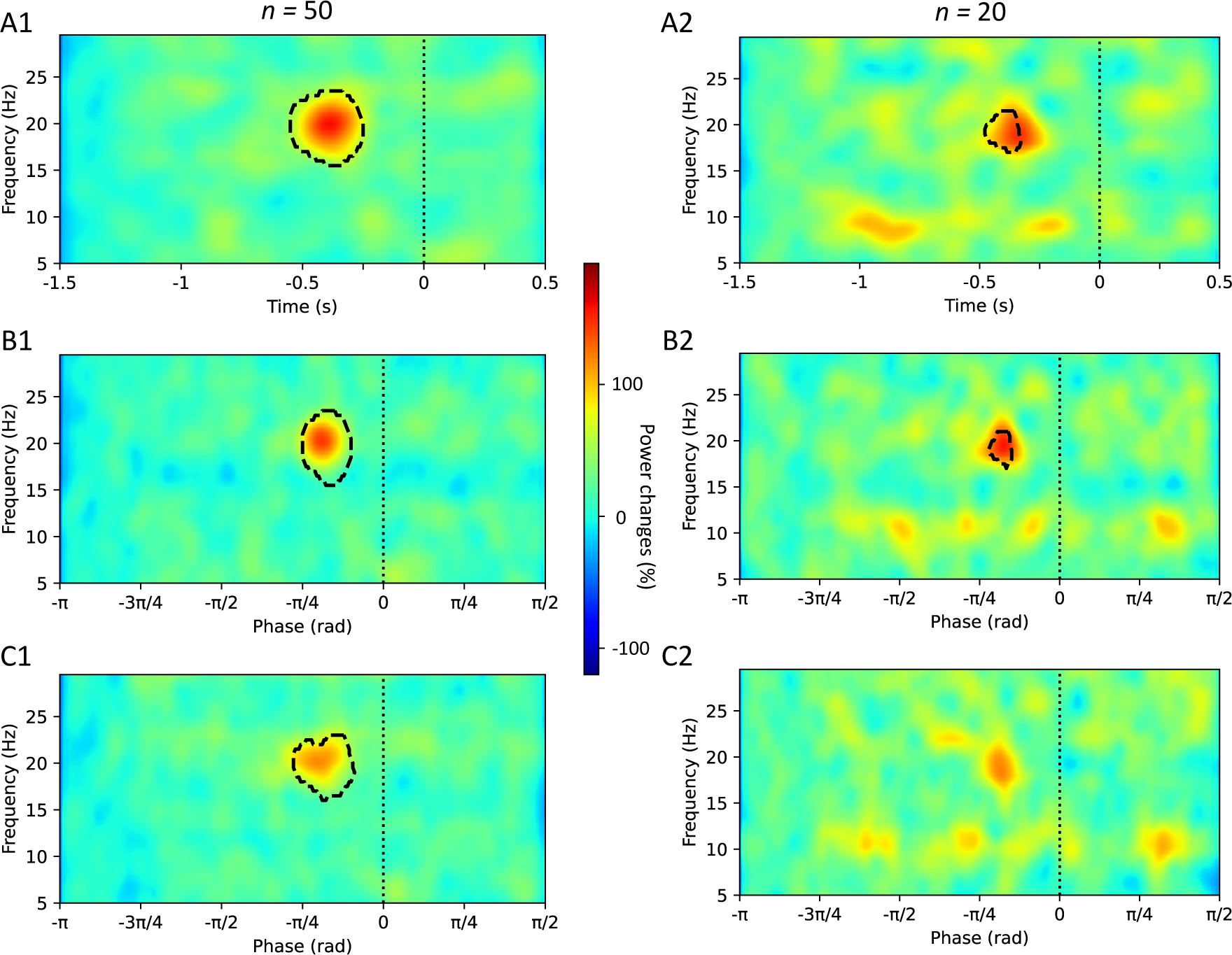
Cycle-frequency maps derived from synthetic EEG generated by the time-locked model. Rows A, B and C illustrate different power representations: A1 shows the map obtained by method A (conventional time-frequency transformation of fixed-length segments, ***X***^(2)^). B1 presents the map computed by method B (cycle-frequency transformation employing segments with a fixed duration based on the mean respiratory rate, ***Y***^(2)^). C1 displays the map obtained by method C (cycle-frequency transformation using cycle-adapted segmentation ***Z***^(2)^). Left plots (A1, B1, C1) depict results from n=50 simulted trials, while right plots (A2, B2, C2) show equivalent representations from another simulation with n=20 trials. The color scale expresses the percentage of power changes with respect to the baseline. Black dotted lines enclose statistically significant areas found by the permutation test.

Finally, using method C, statistically significant activity around the burst was identified only in simulations with n=50 trials (panel C1). In contrast, with only n=20 trials, a small non significant area proximal to the burst was visible in panel C2, which can be potentially confounded with other areas containing spurious activity.

#### Robustness to the number of trials

We generated from *n*=10 to *n*=130 simulated trials to quantify the robustness of T-F and C-F techniques in the context of limited quality data availability. To generate the breathing rythms, we kept the mean breathing rate μ_r_ = 4s and the variance ο_r_ = 1. We used both cycle-locked and time-locked generators and we repeated the procedure 10 times to obtain the mean value of F-statistics and standard errors.

Figure 5-a shows results for cycle-locked potentials. Method C clearly outperforms methods A and B in terms of statistical significance (higher F-statistics) even with as few as 10 trials. This gap in F-statistics widens as the number of trials increases. The phase representation in method C improves the signal-to-noise ratio, leading to better detection. As expected, methods A and B also improve with more trials, but their significance levels remain lower than method C.

**Figure 5:**
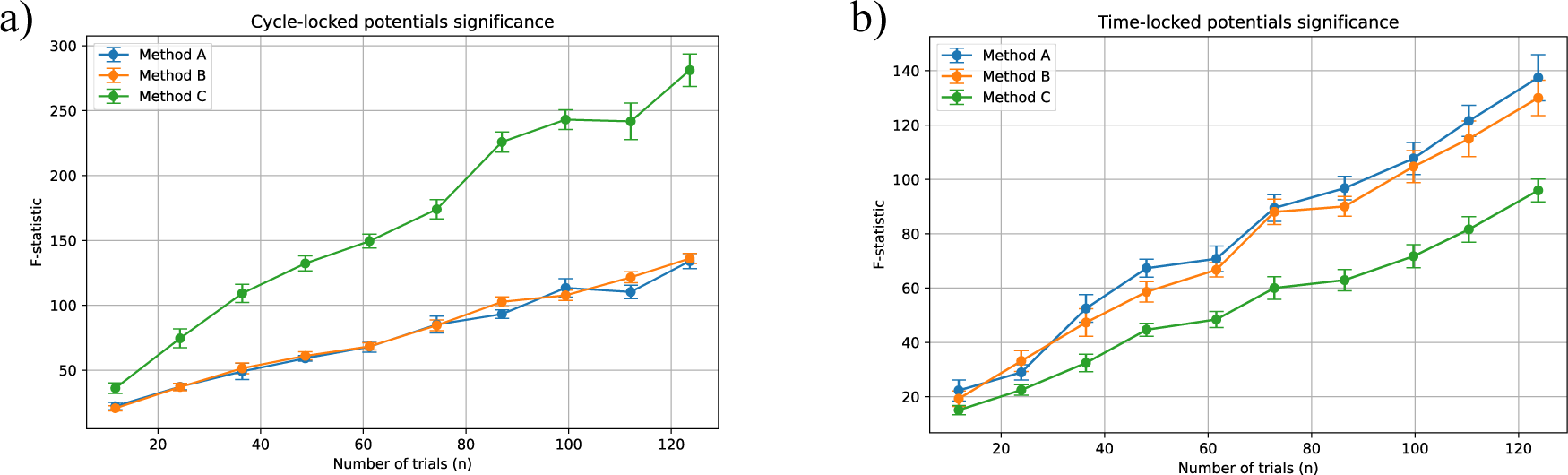
a) Cycle-locked potentials and b) time-locked potentials significance measured with the F-statistic against increasing number of trials and fixed respiratory variability.

Conversely, Figure 5-b displays results for time-locked potentials. Here, methods A and B outperform method C. However, the difference in F-statistics is smaller compared to cycle-locked potentials. Performances of method C could be explained by a leakage effet produced by a short fixed time latencies of potentials relative to inspiration triggers.

Regarding the three methods examined, conventional time-frequency representations are effective for detecting time-locked activity, while cycle-frequency representations are better suited for cycle-locked activity, particularly when a sufficient number of trials ensures a good signal-to-noise ratio. However, with lower signal-to-noise ratios (n ≤ 20 trials), methods A and B may struggle to accurately identify phase-locked activity. Similarly, method C cannot correctly identify time-locked activity with a reduced number of trials.

*Robustness to inter-trial variability*.

To assess the robustness of the different methods to breathing variability, we performed an ensemble of simulations of n=50 trials with μ_r_ = 4s and increasing ο_r_ from 0.2 to 2.2 in steps of 0.2. We repeated 10 times the procedure to obtain mean values of F-statistic and its standard errors.

In Figure 6-a, we can observe that when breathing is almost constant (ο_r_ = 0.2), the three methods exhibit similar high performances in detecting cycle-locked potentials. However, method C proves to be the most robust when respiratory cycles are highly variable. The opposite is observed for time-locked activties, as illustrated in Figure 6-b. Methods A and B perform better than a cycle-based segmentation. This is due to the fixed time latency of potentials relative to the inspiration triggers, which in the case of long breaths, causes a leakage effect in method C. Consequently, the power of the burst is distributed over a wider area in the maps, reducing its power level and thus providing less pronounced significance.

**Figure 6:**
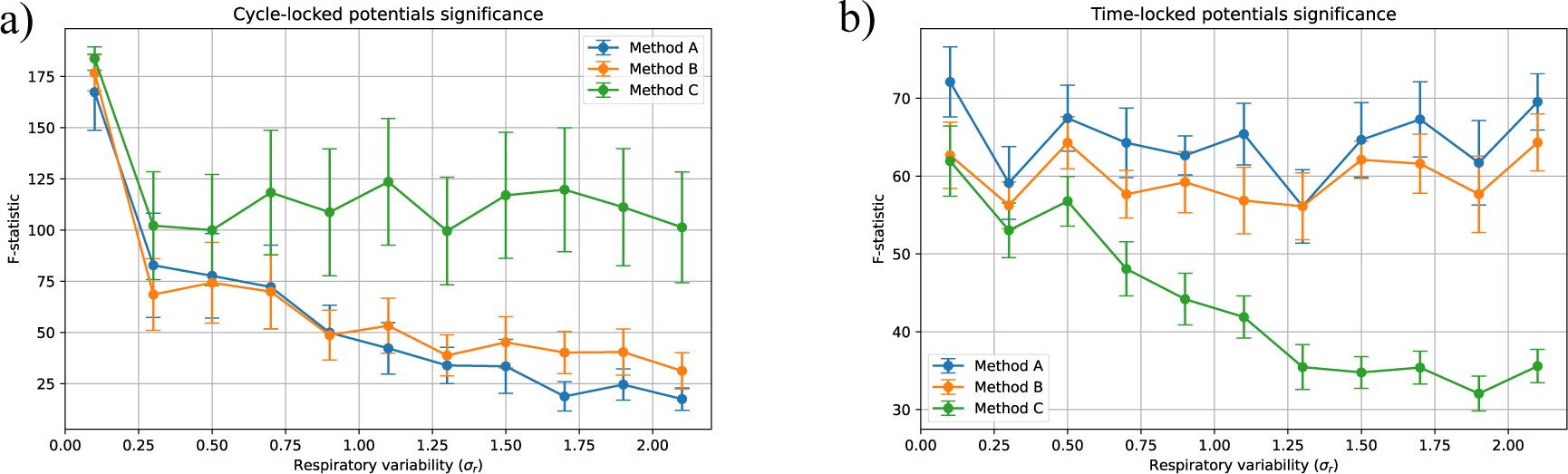
a) Cycle-locked potentials and b) time-locked potentials significance measured with the F-statistic against increasing respiratory variability and fixed number of trials.

Method B is more robust to respiratory variability than method A when considering cycle-locked potentials. Additionally, method B outperforms method C when considering time-locked potentials. Therefore, method B could serve as a suitable compromise when the presence of cycle-or time-locked potentials is uncertain.

### Tests on real data

To illustrate the potential of our proposal, we compared qualitatively standard Time-Frequency (T-F) and Cycle-Frequency (C-F) maps using real data from two subjects with different breathing patterns (see Table 1; see maps in Figures A1 and A2 from the appendix). Subject 1 had the fastest breathing rate in UB, the slowest rate in LB and more overlapped trials using fixed segmentation parameters (*t_pre_* = −1.5 s; *t_post_* = 0.5 s).

**Table 1:** Mean respiratory periods 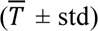 and number of trials *n* (overlapping segments).

|  | Unloaded breathing |  | Loaded breathing |  |
| --- | --- | --- | --- | --- |
| | $\bar{T}_{UB} \pm \text{std}$ | $n$ (overlapping) | $\bar{T}_{LB} \pm \text{std}$ | $n$ (overlapping) |
| Subject 1 | $2.72 \pm 0.47$ s | 220 (54) | $8.73 \pm 2.01$ s | 66 (0) |
| Subject 2 | $3.34 \pm 0.36$ s | 180 (1) | $5.59 \pm 0.99$ s | 108 (0) |

As for the tests on synthetic data, we examined the statistical differences in pre-inspiratory activity relative to the baseline using permutation tests for both subjects and both conditions separately. Power maps for subject 1 are presented in the Appendix (Figures A1 and A2). Of note, due to the longer duration of breathing cycles in LB, C-F maps display an extended temporal window, causing the potentials to appear compressed compared to their counterparts in T-F maps (see Figure A2 in the Appendix, rows B and C). Beyond this visual effect, longer segments in the cycle-frequency transformation can yield unrealistic values for the onset and duration of respiratory-related potentials. For instance, if we consider that pre-inspiratory potentials start at phase –π/4, the corresponding time for a long breath of T=15 s (tpre=-7.5 s; tpost= 3.25 s) is −1.875 seconds. Although this latency might be plausible according to prior research on slow pre-inspiratory cortical potentials (Hudson et al., 2018; Macefield & Gandevia, 1991; Raux et al., 2007), it is unrealistic for faster activity such as the mu rhythm (Kilavik et al., 2013; Pfurtscheller & Lopes da Silva, 1999). To address this issue, we propose to apply an arbitrary threshold during segmentation. In the LB condition analysed here, we set 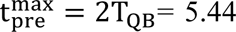 seconds. However, even after applying this threshold, differences in the computed maps did not result in statistically significant outcomes in permutation tests when comparing the three methods (see figures A3 and A4 in the Appendix for more details).

Next, we computed the relative power changes in mu and beta bands with respect to the baseline for each condition and for each method. As shown in Figure 7, the power distributions of both bands across 21 EEG channels differ for the two subjects. However, the average power increase for mu and beta bands was less in the loaded breathing condition compared to unloaded breathing. The decreases in beta power were not significant using the traditional time-frequency analysis (method A) for both subjects when comparing UB and LB conditions. Subject 1 had the most variable breathing, especially in the LB condition; which could explain the lack of significance in mu power. Subject 2 had less variable breathing, which presumably resulted in clearer reductions in power in both mu and beta bands.

**Figure 7:**
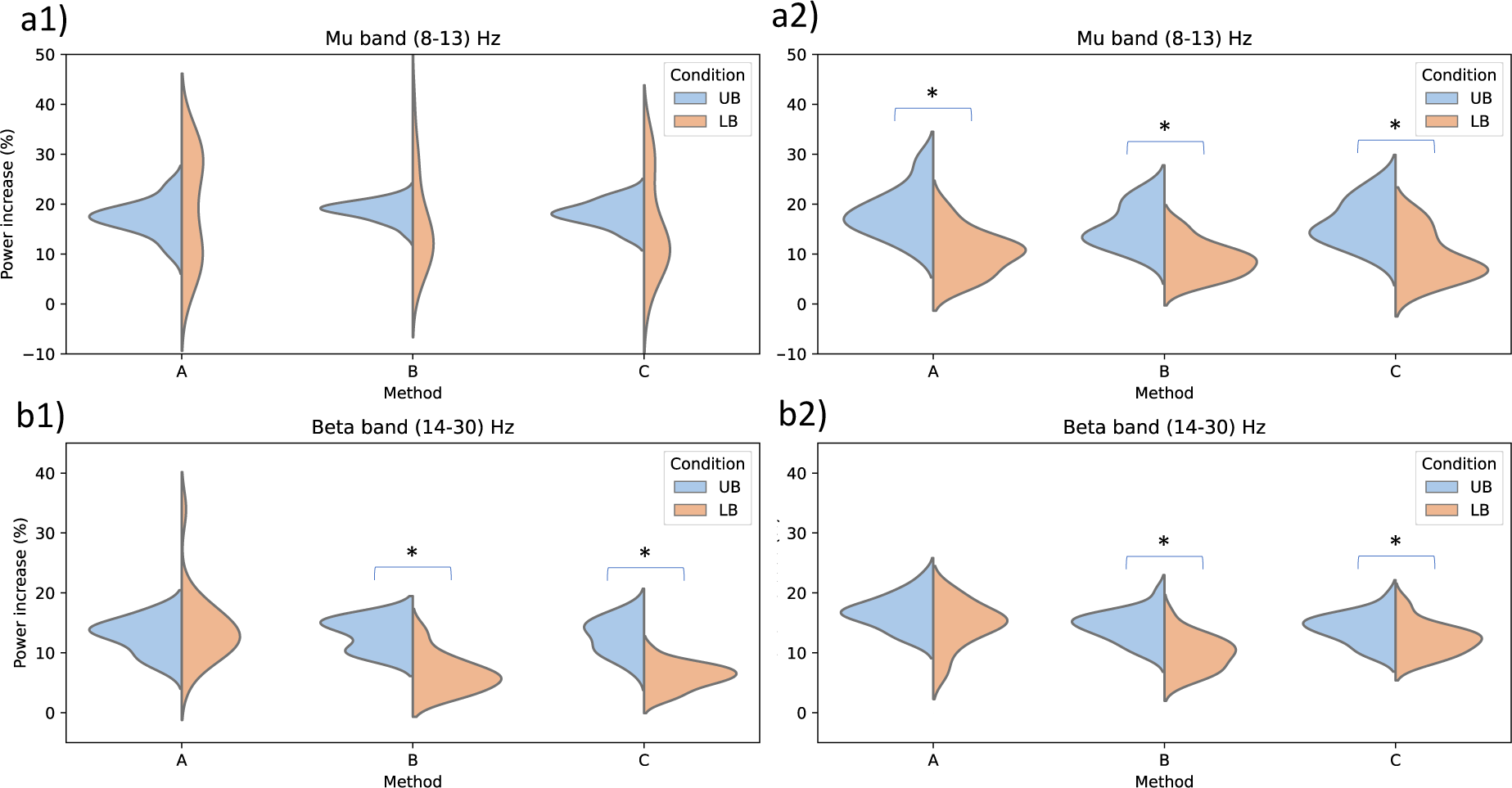
Relative power changes of mu and beta bands across 21 EEG channels in two subjects, comparing respiratory conditions and methods (A: time-frequency transformation, B: cycle-frequency using cycle-averaged segments and C: cycle-frequency using cycle-adapted segments). Left panels correspond to mu (a1) and beta (a2) bands for subject 1. Right panels correspond to mu (a2) and beta bands for subject 2. Asterisks indicate significant differences comparing UB and LB found by Wilcoxon signed-rank test at p-value < 0.01.

Methods pairwise comparisons were conducted on the power distributions using nonparametric signed rank tests. In subject 1, significant differences were observed only when comparing A versus B and A versus C for beta power in the LB condition. In contrast, in subject 2, power distributions were significantly different in all cases except for mu power in the LB condition.

## DISCUSSION

In this study, we introduced a new procedure for analyzing event-related spectral perturbation, called cycle-frequency (C-F) transformation, specifically designed to examine cycle-locked events by adjusting trial lengths to match the duration of each cycle. Non-strictly-periodic EEG activities are prevalent in many paradigms. To address this variability, several studies have employed time-warping methods to enhance the characterization of temporal dynamics in neural activity. This technique, previously applied in analyses of self-paced rhythmic movements (Chemin et al., 2018) and gait analysis (J. Wagner et al., 2019), involves the contraction or dilation of temporal signals to align trial lengths with a specified mean period. The C-F transformation offers an alternative technique where trial segmentation is adapted to the duration of each breath.

We have demonstrated that, for the respiratory context, the use of C-F representation has several advantages over the conventional time-frequency (T-F) procedure. One of the advantages is the ability to normalize the time relative to the phase of the cyclic activity. As the duration of each breath may be highly variable (Fiamma et al., 2007), this normalization facilitates the averaging of trials and their comparison across different conditions and subjects. Another advantage is the possibility to set the latency and duration of the baseline adapted to the cycle length. This baseline placement strategy systematically avoids the overlap of the reference period with active periods of neighbouring events.

As a result, the analysis of respiratory-related cortical activity becomes more robust to inter-trial variability (due for instance to changing states during an experiment). Considering interventions to relieve dyspnea in mechanically ventilated patients, breathing rates are likely to change substantially (Raux et al., 2019). Therefore, our proposal can provide, for example, a benchmark for comparing intervention-related cortical activities during fast and slow breathing.

To evaluate the effectiveness of C-F analysis, we used artificial signals mimicking cycle-locked and time-locked pre-inspiratory EEG potentials. Our findings demonstrate that C-F analysis enhances the detection of cycle-locked potentials compared to conventional T-F procedures, which may obscure cycle-locked activity due to temporal misalignment during the averaging process. Conversely, while time-locked acivity can be efectively detected by T-F maps, C-F analysis fails when signal-to-noise levels were lower (n < 20 trials). Employing cycle-frequency maps with segments of fixed duration based on the mean respiratory rate may offer a compromise for detecting both time- and cycle-locked activities.

To illustrate our approach with real data, we applied both cycle-frequency and time-frequency analyses to EEG data from two subjects breathing normally (UB) and breathing through a respiratory load (LB). Cycle-frequency methods may reveal respiratory-related activity that was not evident with the traditional method as shown here, and in our previous studies using time-frequency analyses of loaded breathing (Hudson et al., 2016) and hypocapnic breathing (Dubois et al., 2016). In contrast with tests on synthetic data, the analysis of pre-inspiratory potentials compared to the baseline in real EEG did not yield statistical significant results. The statistical test, designed for differences in pre-inspiratory activity compared to baseline in the synthetic data, is a very restrictive model for real-world data with potential confounding factors such as measurement noise and subject-specific fluctuations. In addition, individual sensations and capacity to compensate for breathing constraints are highly subject-specific. We presented results from two subjects, making difficult the generalization of findings, specially in respiratory experiments.

Despite the mitigated results on real data, analyzing average power across channels and focusing on specific frequencies and latencies revealed interesting results. Consistent with similar movement preparation paradigms, we expected a decrease in mu and beta power in the LB condition compared to UB. Paired statistical tests showed a significant decrease in beta power for both subjects in the LB condition using cycle-frequency methods, while the traditional time-frequency method revealed no significant effects.

These results partially validate our hypothesis of cycle-locked potentials, but further research is needed to determine the extent to which time- and cycle-locked activities coexist within the same experimental condition. As suggested by tests using synthetic data, time-locked and cycle-locked activity can significantly impact the reliability of power representations. In this regard, using both time-frequency (T-F) and cycle-frequency (C-F) maps holds promise for studying overall cortical activity related to breathing.

By analyzing real data we also conclude that fast cycle-locked potentials might not start with long latencies in very long cycles. While there is evidence that readiness potentials may commence, under certain conditions, as early as 2 seconds before the onset of the movement (Hudson et al., 2018; Macefield & Gandevia, 1991; Raux et al., 2007), such latency is unrealistic for faster rhythms. Therefore, we proposed implementing a threshold to restrict the maximum length of EEG segments to be converted into the cycle domain, thereby avoiding time compression that could blur fast transient activity. This threshold should be a reasonable value based on the available evidence, such as the upper quartile of breathing periods at rest or twice the mean period.

The proposed method presents some limitations that should be carefully considered. Since the downsampling of C-F maps leads to a lower temporal resolution of high frequencies, it is crucial to intially evaluate whether parameters *l_min_* and *f_max_* accurately represent the desired frequencies. In datasets containing trials with significant variations in duration, it may be necessary to exclude the shortest trials to raise *l_min_* and therefore increase *f_max_*. We also notice that the downsampling procedure in the CF transformation may have an impact on the statitistical significance of permutation tests (reducing the number of pixels can dilute the power of specific, time-located activities).

Another area for future investigation concerns the influence of cycle asymmetry, where inspiration and expiration durations may differ significantly. Our current model for generating respiratory markers focused solely on total cycle duration, following a log-normal distribution for simplicity. However, incorporating more detailed and complex rhythm generators, as proposed by Alian et al. (Alian et al., 2022), could provide more realistic breathing patterns. This would allow for separate analyses of brain activity specifically evoked by inspiration and exhalation.

Finally, future works should explore the generalizability of C-F analysis to other types of self-paced responses to investigate the potential of this technique for the study of cycle-locked potentials. This research could have notable implications for the characterization of respiratory and limb movement disorders and lead to a better understanding of the neural control of rhythmic movements.

## CONCLUSION

The objective of this paper is to introduce a new tool for computing event-related spectral perturbations in cortical activity associated with cycle or internal-paced movements. This tool is intended to be used in conjunction with conventional time-frequency (T-F) analysis rather than as a replacement, as time-locked potentials may coexist with cycle-locked potentials. Supplementary material in https://doi.org/10.17605/OSF.IO/ZM5UG includes the code and sample data to produce C-F maps to enable the reproduction of our results and facilitate further investigations.

## APPENDIX

**Table A1:** List of parameter names, values used and justification for these values.

| Parameter | Name | Value | Remarks |
| --- | --- | --- | --- |
| $\eta$ | Number of cycles | 3 to 12 | Increasing value with increasing wavelet frequencies (from 2.5 to 14.75 Hz). |
| $l_{min}$ | Shortest breath | 2 s (synthetic);<br>1.5 s (real) | The minimal trial duration determines the lower analysis frequency, $f_{min}$. |
| $f_{min}$ | Minimal frequency | 5 Hz | Minimun frequency of interest (lower value in power maps).<br>Must satisfy $f_{min} \geq 3/l_{min}$ to ensure good frequency resolution. |
| $f_{max}$ | Maximal frequency | 30 Hz | Maximal frequency of interest (upper value in power maps).<br>Must satisfy $f_{max} \leq f_s/2$ to follow Nyquist criterion. |
| $f_b$ | Frequency bin | 0.5 Hz | Frequency step from $f_{min}$ to $f_{max}$ |
| q | Number of frequencies | 50 | Determines the number of vertical pixels of the power maps. |
| $\phi_s$ | Start phase | $-\pi$ rad ($-180^\circ$) | Inclusion of half of previous breath (start of previous expiration) to study pre-inspiratory activity. |
| $\phi_e$ | End phase | $\pi/2$ ($90^\circ$) | Inclusion of half inspiration. |
| $\Delta\phi$ | Phase interval of interest | $3\pi/2$ ($270^\circ$) | $\Delta\phi = \phi_e - \phi_s$ determines the phase span of the power maps. |
| $r_\phi$ | Phase bin | $1^\circ$ | Unit value provides a trade-of between horizontal resolution and computational time. |
| p | number of phase values (bins) | 270 | Must satisfy $p \geq \frac{2f_{max}}{l_{min}}$<br>Determines the number of horizontal pixels of power maps. |
| $k_i$ | Trial-dependent Downsampling factor | $p/m_i$ | Variable factor to scale from time domain to phase domain. Depends on p and the number of samples per segment ($m_i$). |

### Analysis of real EEG in motor-related cortical areas

We performed time-frequency and cycle-frequency analyses using the data from the two subjects. After obtaining different maps for each channel, we computed an average map covering the pre-motor area (average of positions F3, Fz, F4, FC1, and FC2) and an average map covering somatosensory positions (average of C3, Cz, C4, CP1, and CP2) to investigate differences between methods A, B and C. These areas are of particular interest in the analysis of respiratory-related potentials as they reflect the preparation of breaths and the sensory perception of breathing. For both subjects, permutation tests did not identify statistically significant changes with respect to the baseline period. Figures A1 and A2 illustrate Subject 1 averaged maps for pre-motor and somatosensory areas in UB and LB conditions, respectively.

**Figure A1:**
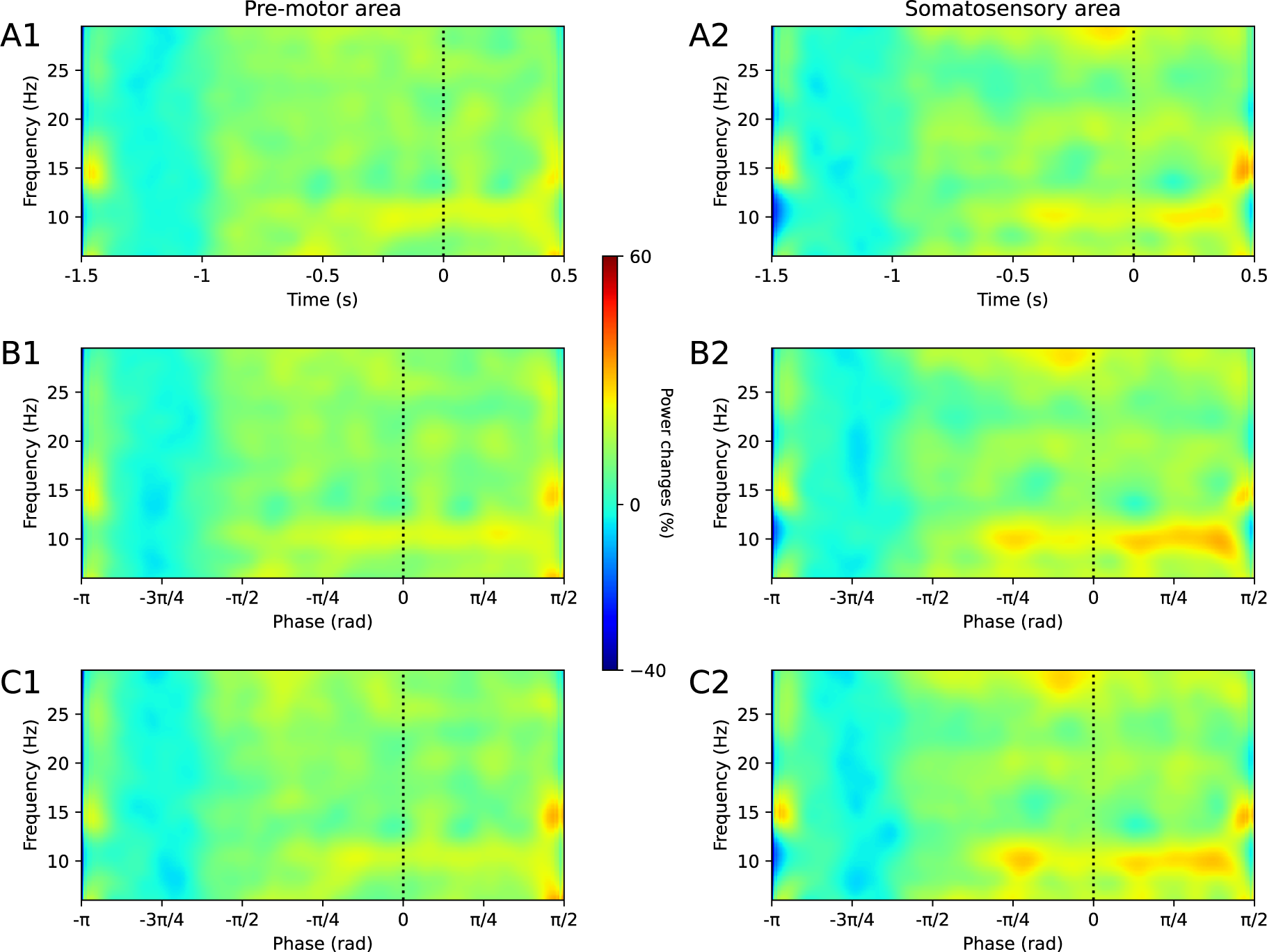
Power maps obtained from real EEG during unloaded breathing (n=166 trials). The left column (A1, B1, C1) displays average maps using electrodes in the pre-motor area, while the right column (A2, B2, C2) presents average maps from the somatosensory area (see Results for EEG positions). The top row (A1, A2) illustrates conventional time-frequency maps using fixed segmentation ([−1.5; 0.5] sec); the middle row (B1, B2) showcases cycle-frequency maps with fixed segmentation based on the average breathing period; and the bottom row (C1, C2) displays cycle-frequency maps with cycle-adaptive segmentation.

**Figure A2:**
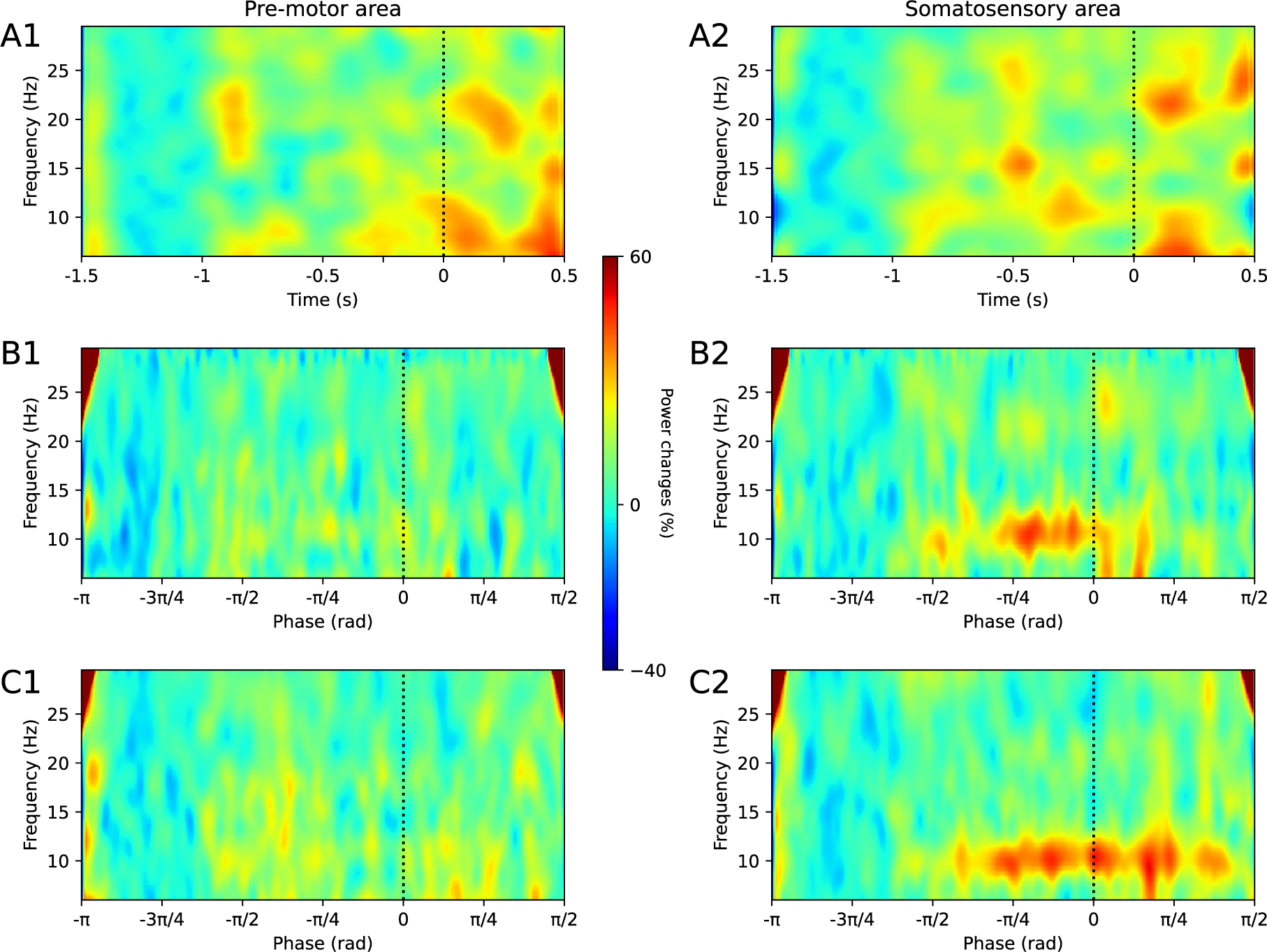
Power Maps Derived from Real EEG during loaded breathing (n=66 trials). The left column (A1, B1, C1) displays average time-frequency maps utilizing electrodes in the pre-motor area, while the right column (A2, B2, C2) presents average maps from the somatosensory area. The first row (A1, A2) illustrates conventional time-frequency maps employing fixed segmentation ([−1.5; 0.5] sec); the second row (B1, B2) showcases cycle-frequency maps with fixed segmentation based on the average breathing period; and the third column (C1, C2) exhibits cycle-frequency maps utilizing cycle-adaptive segmentation.

We also computed the relative power for each map by dividing the mean power value during the pre-inspiratory latencies of interest ([-π/4, 0]) by the baseline period (20% of the entire pre-inspiratory time), within the frequencies [9-13] Hz for the mu band (see Figure A3) and [14-30] Hz for the beta band (see Figure A4). All power values were approximately normally distributed, and t-tests revealed no significant differences when comparing between methods and conditions.

**Figure A3:**
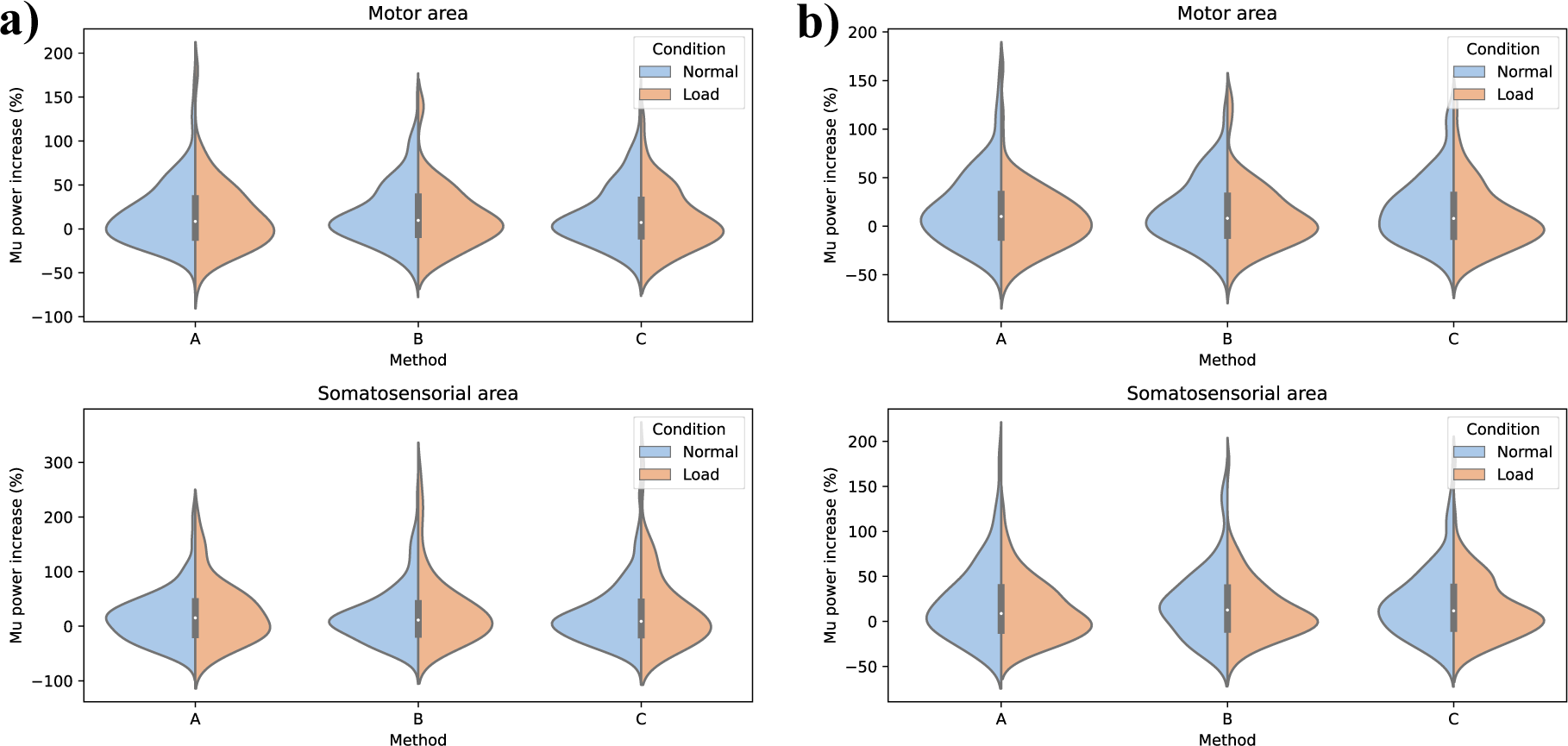
Relative power in the mu band (motor and somatosensory areas) obtained by methods A, B, and C in UB and LB conditions. Panel a) visualizes power values for Subject 1, while panel b) shows power values for Subject 2.

**Figure A4:**
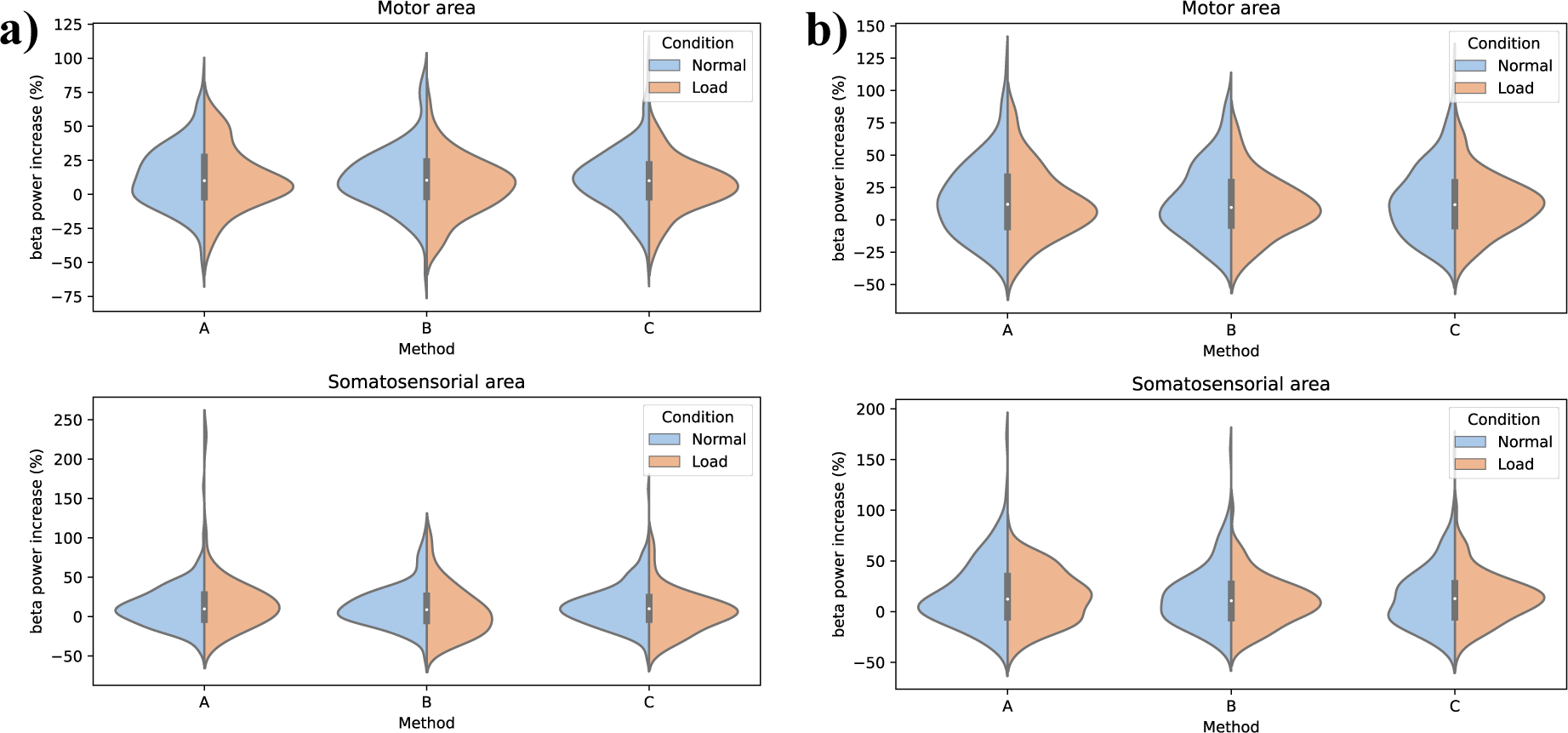
Relative power in the beta band (motor and somatosensory areas) obtained by methods A, B, and C in UB and LB conditions. Panel a) visualizes power values for Subject 1, while panel b) shows power values for Subject 2.

